# SAFE-clustering: Single-cell Aggregated (From Ensemble) Clustering for Single-cell RNA-seq Data

**DOI:** 10.1101/215723

**Authors:** Yuchen Yang, Ruth Huh, Houston W. Culpepper, Yuan Lin, Michael I. Love, Yun Li

## Abstract

**Motivation:** Accurately clustering cell types from a mass of heterogeneous cells is a crucial first step for the analysis of single-cell RNA-seq (scRNA-Seq) data. Although several methods have been recently developed, they utilize different characteristics of data and yield varying results in terms of both the number of clusters and actual cluster assignments.

**Results:** Here, we present SAFE-clustering, Single-cell Aggregated (From Ensemble) clustering, a flexible, accurate and robust method for clustering scRNA-Seq data. SAFE-clustering takes as input, results from multiple clustering methods, to build one consensus solution. SAFE-clustering currently embeds four state-of-the-art methods, SC3, CIDR, Seurat and t-SNE + *k*-means; and ensembles solutions from these four methods using three hypergraph-based partitioning algorithms. Extensive assessment across 12 datasets with the number of clusters ranging from 3 to 14, and the number of single cells ranging from 49 to 32,695 showcases the advantages of SAFE-clustering in terms of both cluster number (18.2 - 58.1% reduction in absolute deviation to the truth) and cluster assignment (on average 36.0% improvement, and up to 18.5% over the best of the four methods, measured by adjusted rand index). Moreover, SAFE-clustering is computationally efficient to accommodate large datasets, taking <10 minutes to process 28,733 cells.

**Availability and implementation:** SAFEclustering, including source codes and tutorial, is freely available at https://github.com/yycunc/SAFEclustering.

**Supplementary information:** Supplementary data are available at *Bioinformatics* online.

## 1 INTRODUCTION

RNA sequencing (RNA-seq) has been widely used to study gene regulatory networks underlying the complex processes of cellular proliferation, differentiation and reprograming (Trapnell *et al*., 2014; Treutlein *et al*., 2014; Darmanis *et al*., 2015). However, for most genes, their expression levels are found to vary dramatically across cell types and in different individual cells (Tang *et al*., 2010; Buganim *et al*., 2012; Shalek *et al*., 2013). Therefore, bulk RNA-seq, measuring the average expression across many cells of different cell types, may mask the real functional capacities of each cell type (Trapnell *et al*., 2014). Comparatively, single-cell RNA sequencing (scRNA-Seq) enables researchers to investigate the cellular heterogeneity in gene expression profiles, as well as to determine cell types and predict cell fates, thus presenting enormous potential for cell biology and clinical applications (Treutlein *et al*., 2014; Kalisky and Quake, 2011; Arsenio *et al*., 2014; Jaitin *et al*., 2014; Mahata *et al*., 2014; Grün *et al*., 2015; Jia *et al*., 2017).

Single-cell clustering provides intuitive identification and characterization of cell types from a mass of heterogeneous cells, which can itself be of interest (Rozenblatt-Rosen *et al*., 2017), and can be used as covariates in downstream differential expression analysis (Sun *et al*., 2017; Zhu *et al*., 2017). Because of the importance of clustering for scRNA-Seq data, recently, several algorithms have been developed, including t-Distributed Stochastic Neighbor Embedding algorithm (t-SNE) (Van der Maaten and Hinton, 2008) followed by *k*-means clustering (Grün *et al*., 2015; Shin *et al*., 2015), Seurat (Satija *et al*., 2015), DIMM-SC (Sun *et al*., 2017), SIMLR (Wang *et al*., 2017), SC3 (Kiselev *et al*., 2017), DendroSplit (Zhang *et al*., 2018) and SCANPY (Wolf *et al*., 2018). However, none of the clustering algorithms is an apparent all-time winner across all datasets (Freytag *et al*., 2017). Discrepancies across methods occur both in the estimated number of clusters and in actual single-cell-level cluster assignment. These discrepancies are mainly due to the use of different characteristics of scRNA-Seq data by different methods, for example, different sets of genes used for downstream clustering from different choices of gene level filtering subsetting of gents, transformation and dimension reduction. Individual clustering methods may fail to reveal the true clustering behind a heterogeneous mass (of single cells in this case) when assumptions underlying the methods are violated. Therefore, it is highly challenging, if not impossible, to choose an optimal algorithm for clustering scRNA-Seq data when no prior knowledge on cell types and/or cell type specific expression signatures are given.

In the absence of one single optimal clustering method, cluster ensemble provides an elegant solution by combining results from multiple individual methods into one consensus result (Strehl and Ghosh, 2002; Ghosh and Acharya, 2011). Compared to individual solutions, ensemble methods exhibit two major advantages. First, ensemble improves clustering quality and robustness, as demonstrated in other contexts including analysis of cell signalling dynamics and protein folding (Kuepfer *et al*., 2007; Hubner *et al*., 2005). Second, ensemble methods enable model selection. For example, we and others (Lin *et al*., 2017; Kiselev *et al*., 2017; Freytag *et al*., 2017) observe, in certain datasets, dramatically different estimates for the number of clusters across individual solutions. It is hard to decide on one single solution without any external knowledge or constraints. Cluster ensemble is able to estimate an optimal number of clusters by quantifying the shared information between the final consensus solution and individual solutions (Ghosh and Acharya, 2011). Although the majority may not always be the most accurate in every case and for every cell, a consensus approach tends to outperform each individual method when the optimal method is not known in advance. However, to date, there is no published cluster ensemble approach across multiple types of clustering methods specifically designed for scRNA-Seq data.

To bridge the gap, we have developed SAFE-clustering, <u>S</u>ingle-cell <u>A</u>ggregated (<u>F</u>rom <u>E</u>nsemble) <u>clustering</u>, to provide more stable, robust and accurate clustering for scRNA-Seq data. In the current implementation, SAFE-clustering first performs independent clustering using four state-of-the-art methods, SC3, CIDR, Seurat and t-SNE + *k*-means, and then combines the four individual solutions into one consolidated solution using one of three hypergraph partitioning algorithms: hypergraph partitioning algorithm (HGPA), metaclustering algorithm (MCLA) and cluster-based similarity partitioning algorithm (CSPA) (Strehl and Ghosh, 2002).

## 2 MATERIALS AND METHODS

### 2.1 Overview of SAFE-clustering

Our SAFE-clustering leverages hypergraph partitioning methods to ensemble results from multiple individual clustering methods. The current SAFE-clustering implementation embeds four clustering methods: SC3, Seurat, t-SNE + *k*-means, and CIDR. Fig. 1 shows the overview of our SAFE-clustering method.

**Fig. 1.**
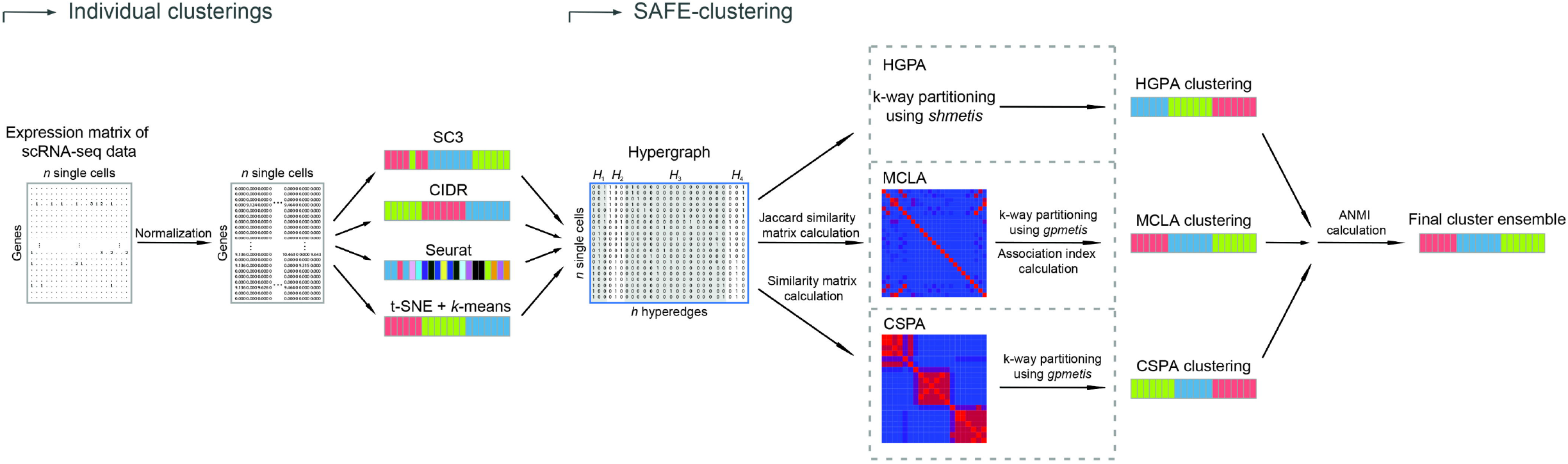
Overview of SAFE-clustering. Log-transformed expression matrix of scRNA-Seq data are first clustered using four state-of-the-art methods, SC3, CIDR, Seurat and t-SNE + *k*-means; and then individual solutions are combined using one of the three hypergraph-based partitioning algorithms: hypergraph partitioning algorithm (HGPA), meta-cluster algorithm (MCLA) and cluster-based similarity partitioning algorithm (CSPA) to produce consensus clustering.

#### 2.2 Expression matrix normalization

SAFE-clustering takes an expression matrix as input, where each column represents one single cell and each row corresponds to one gene or transcript. To make the data well-suited for all four individual clustering methods, Fragments/Reads Per Kilobase per Million mapped reads (FPKM/RPKM) data are converted into Transcripts Per Million (TPM); and UMI counts are converted into Counts Per Million mapped reads (CPM). For CIDR, SC3 and t-SNE + *k*-means, the input expression matrix is log-transformed after adding ones (to avoid taking log of zeros).

#### 2.3 Clustering using four state-of-the-art methods

##### CIDR

To deal with the dropout events in scRNA-seq data, CIDR first identifies dropout candidates from the expression matrix and performs implicitly imputation to mitigate the impact of lowly expressed genes (Lin *et al*., 2017). Then, dissimilarity matrix (Euclidean distance) is calculated between single cells using the imputed data. As CIDR performs principal coordinate analysis (PCoA) to reduce dimensionality, the number of principal coordinates (PCo’s) identified, representing the estimated data dimensionality, heavily influences the final clustering results. Here, the number of PCo’s is determined by the internal *nPC* function, default choice in CIDR. Alternatively, users can visually decide on an ideal number of PCo’s by selecting a threshold at a clear elbow from plotting the proportions of variations explained by the PCo’s (also generated by the *nPC* function). With the selected PCo’s, single cells are hierarchically clustered into 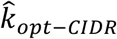 clusters, with 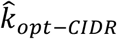 estimated using the Calinski-Harabasz Index (Caliński and Harabasz, 1974).

##### SC3

SC3 adopts consensus clustering, and summarizes the probability of each pair of cells is from the same cluster (Kiselev *et al*., 2017). Quality control (QC) metrics are calculated on the input expression matrix to detect potentially problematic genes and/or single cells. Although gene-level filtering is recommended by SC3, for 9 out the 12 benchmarking datasets, all genes would be filtered out and clustering cannot be performed. Therefore, we set the gene filtering option to be “FALSE”. In order to speed up computation, we first use the Tracy-Widom method (Tracy and Widom, 1994; Patterson *et al*., 2006) to estimate the number of clusters, denoted by 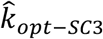. With the estimated 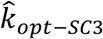, matrices of Euclidean, Pearson and Spearman (dis)similarity metrics are calculated among single cells, followed by *k*-means clustering. Based on A-means results across the three different (dis)similarity matrices, a consensus matrix is computed using CSPA, followed by a hierarchical clustering to assign the single cells into 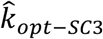 clusters.

For the two PBMC mixture datasets (both with >5,000 single cells), via SC3 default implementation, support vector machines (SVM) is employed to further speed up computation. Specifically, a subset of single cells is randomly selected to form the training dataset where a SVM model with a linear kernel is constructed, using the *svm* function in R-package e1071. The default minimum number of single cells to run SVM is set to be 5,000 (SC3 option *svm_max*, default = 5,000). The trained SVM is then used to predict the cluster labels of the remaining single cells.

##### Seurat

Seurat embeds an unsupervised clustering algorithm, combining dimension reduction with graph-based partitioning methods. First, expression matrix is filtered to remove genes expressed in <3 single cells and single cells with <200 expressed genes. Then, the expression data of each single cell is scaled to a total of 10,000 molecules and log-transformed following the procedure described in Macosko *et al*. (2015) (Macosko *et al*., 2015). After that, undesired sources of variations are regressed out. Single cells with <200 expressed genes would be considered as “NA” in the final Seurat clustering results. Data dimensionality is reduced via principal component analysis (PCA) with the principal components (PCs) selected by the *nPC* function in the CIDR package. Graph-based clustering is carried out using the smart local moving algorithm (SLM) (Waltman and van Eck, 2013) with the *resolution* parameter set to be 0.9. For small datasets, Seurat has been reported not to work well (Waltman and van Eck, 2013) and has a tendency to assign all single cells into one cluster when the *resolution* parameter is set to be 0.9. We therefore increase the *resolution* parameter from 0.9 to 1.2 when the number of single cells is less than 200.

##### t-SNE + k-means

t-SNE followed by *k*-means clustering is a popular method for single cell clustering, where high dimensional data are first reduced into a lower dimensional subspace by t-SNE algorithm and then the lower-dimensional data are clustered with *k*-means. Here, we use the Rtsne package with default parameters to reduce normalized expression data into three dimensions by default (Users can specify otherwise via option *dims*, detailed in **Supplementary Results**). However, when the number of input single cells is small, users may run into the problem that the default *perplexity* of 30 is too big, for example, for small datasets. Since t-SNE has been shown to be reasonably robust across *perplexity* values ranging from 5 to 50 (Van der Maaten and Hinton, 2008), we set the *perplexity* to be 10 when the input data contain <200 single cells. More evaluations on the perplexity parameter are presented in **Supplementary Results**.

Results from *k*-means clustering can vary dramatically across different runs even with the same input data and same parameters because of random initial cluster centers. To mitigate this potentially highly stochastic behavior, we use the ADPclust R-package (Wang and Xu, 2015) to first estimate the centroids. ADPclust can also estimate the number of clusters. Therefore, in our SAFE-clustering implementation, we perform *k*-means clustering using the centroids and number of clusters estimated through ADPclust.

#### 2.4 Hypergraph Partitioning Cluster Ensemble Algorithms

After obtaining clustering results from different individual methods, we perform cluster ensemble to provide a consensus clustering using one of the three hypergraph-based partitioning algorithms: HGPA, MCLA and CSPA, as described in Strehl and Ghosh (2002). Moreover, certain single cell(s) may be excluded from clustering by some individual clustering method(s) due to quality control filter(s) of the corresponding method(s). Ensemble approach can provide a consolidated assignment for these single cells by borrowing information from solutions of the other methods.

We start with transforming the output labels of each clustering method into a hypergraph. Briefly, for the *j^th^* clustering method, we use *v_ik_* (note subscript *j* is omitted for presentation brevity) to denote the *i^th^* row of the hypergraph *H_j_*, which is the row vector for the cluster labels (coded as binary dummies or indicator functions) of the *i^th^* single cell, where

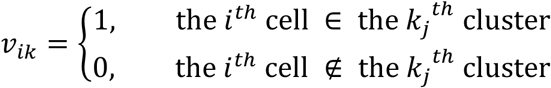

and *k_j_* = 1,2,…, *K_j_*, with *K_j_* being the total number of clusters from the *j^th^* clustering method. Here, each column is a hyperedge, representing one particular cluster identified by that method. An overall hypergraph *H* is constructed by combining individual hypergraphs (from individual methods).

#### HGPA

HGPA directly partitions hypergraphs by cutting a minimal number of hyperedges. We adopt the approach described in Karypis *et al*. (1999), where the authors developed a fast and efficient multilevel hypergraph partitioning algorithm through recursive bisection. Specifically, we perform a fc-way hypergraph partitioning using the *shmetis* program in the hMETIS package v. 1.5 (Karypis *et al*., 1999) for a range of *k* from 2 to max(*K_j_*), *j* = 1,2,3, and 4 for the four different individual clustering methods and *K_j_* again for the total number of clusters from the *j^th^* method. The parameter *UBfactor* is set at 5, so that in any bisection, each of the two partitions contains 45 - 55% of the total number of vertices.

#### MCLA

Unlike HGPA, MCLA starts with computing pairwise Jaccard similarities (*S_J_*) among all the hyperedges. Specifically, for any two hyperedges *h_p_* and *h_q_*:

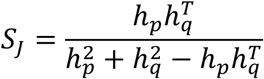

where *p* and *q* = 1,…, *h*, where *h* is the total number of hyperedges, which equals to the sum of estimated cluster numbers from individual solutions. With the calculated similarity matrix, all the hyperedges are partitioned into *k* meta-clusters using the *gpmetis* program in the hypergraph partitioning package METIS v. 5.1.0 (Karypis and Kumar, 1998).

An association index *AI*(*MC_ci_*) is computed to represent the association between meta-cluster *c* and the *i^th^* single cell, by averaging the vertices *v_ch_* of the corresponding hyperedges:

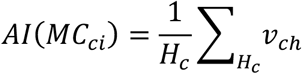

where *h* ∈ *H_c_* is the set of hyperedges assigned in meta-cluster *c*. Each single cell is assigned to the meta-cluster with the highest association index. However, some of the *k* clusters may be empty due to no single cells having the highest association index with the cluster(s) (Strehl and Ghosh, 2002). Under that scenario, we will re-label the single cells into *k*′ clusters, where *k*′ is the number of non-empty clusters.

#### CSPA

CSPA also starts with computing pairwise similarities. In contrast to MCLA, CSPA defines the similarity between two single cells to be 1 if they are *always* assigned to the same cluster, and 0 if they are *never* assigned to the same cluster. The *n*×*n* (where *n* is the number of single cells) similarity matrix *S* can be simply constructed by

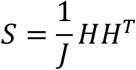

where *H* is the overall hypergraph, and *J* is the total number of individual clustering methods, here *J* = 4. For CSPA, similar to MCLA, we also use the *gpmetis* program in the METIS v. 5.1.0 package.

### 2.5 Performance evaluation using Average Normalized Mutual Information (ANMI)

Since individual methods cluster the single cells into their own optimal number of clusters, we need to estimate an overall optimal cluster number 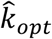 using each of the three ensemble algorithms. For this purpose, we have implemented consensus clustering for a set of *k_e_* = (2,3,…, *K_e_*), where *K_e_* = *max*(*K_j_*) and *j* = 1,2,3 and 4 again for the four individual clustering methods, using each of the three algorithms. We evaluate the performance at each *k_e_* by measuring the shared information between the inferred and true original cluster labels (i.e., mutual information) using the Normalized Mutual Information (NMI) metric, defined in Ghosh and Acharya (2011):

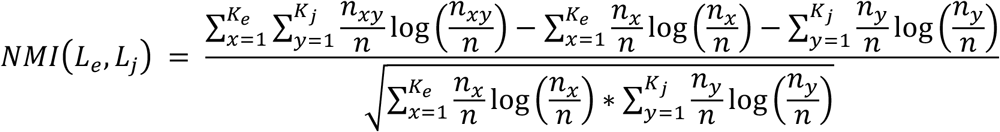

where *L_e_* and *L_j_* are the labels from ensemble and from the *j^th^* method with *K_e_* and *K_j_* clusters, respectively. *n* is the total number of single cells; *n_y_* denotes the number of single cells assigned to a specific cluster *y*(*y* = 1,2,…,*K_j_*) by method *j*; similarly *n_x_* denomtes the number of single cells assigned to cluster *x*(*x* = 1,2,…, *K_e_*) via ensemble; and *n_xy_* represents the number of single cells shared between cluster *y* (from the solution of the *j^th^* individual method) and cluster *x* (from the ensemble solution).

We calculate Average Normalized Mutual Information (*ANMI*) (Strehl and Ghosh, 2002) between each consensus/ensemble solution and each solution from the individual methods. For a particular ensemble solution, the average *ANMI* across individual methods quantifies its similarity to the solutions from individual methods. The ensemble solution with the highest average *ANMI* value (again, average across four individual methods) is selected as the final cluster ensemble 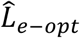 with the estimated cluster number 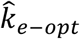:

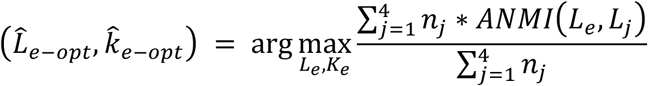

where *n_j_* is the total number of single cells clustered by individual method *j*; and *K_e_* is the number of clusters from an ensemble solution. Note this “average” is more precisely a weighted average rather than a plain average across individual methods unless all methods clustered the same number of single cells (e.g., without removing or failing to cluster any single cell(s), *n_j_* = *n* for all *j*’s). When users simultaneously employ multiple partitioning algorithms (note our default is one single algorithm), the optimal cluster ensemble is given by:

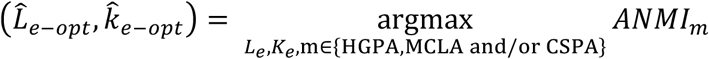

### 2.6 Summary of SAFE-clustering

Run four individual clustering methods and get a *Y*_4×*n*_ matrix of cluster labels. *n* is the total number of single cells.

Construct hypergraph *H* = {*H*_1_, *H*_2_, *H*_3_, *H*_4_}

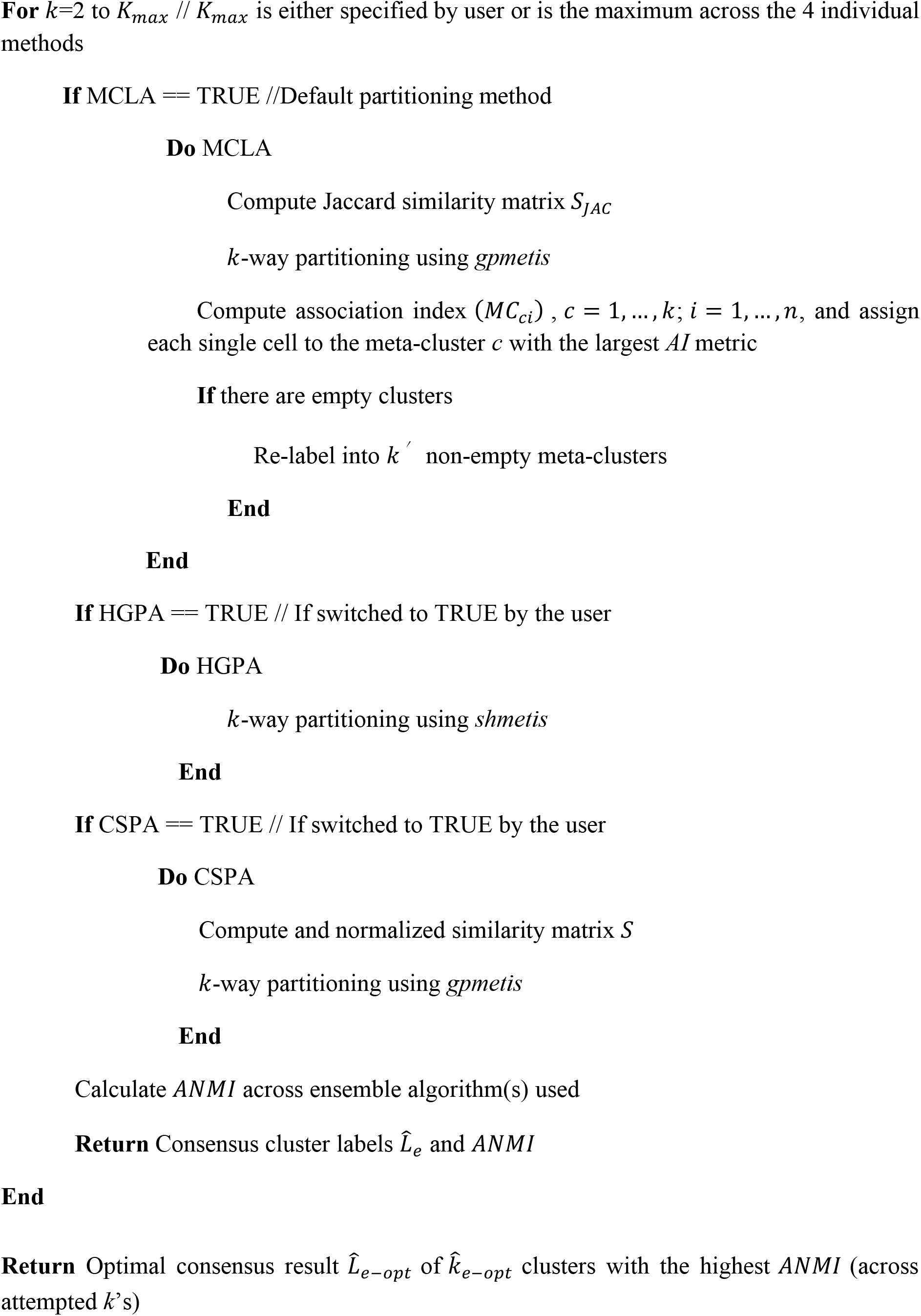

### 2.7 Benchmarking datasets

For performance evaluation, we carried out clustering analysis on 12 benchmark scRNA-Seq datasets (Table 1) (Darmanis *et al*., 2015; Baron *et al*., 2016; Biase *et al*., 2014; Ting *et al*., 2014; Yan *et al*., 2013; Zeisel *et al*., 2015; Zheng *et al*., 2017), using our SAFE-clustering and the four individual clustering methods. All these datasets have pre-defined gold/silver-standard (we call “true”) cell type information. We used default parameters for 10 out of the 12 datasets, with the two exceptions being the 2 PBMC mixture datasets (each with >28,000 single cells). For SC3, gene-level filtering option was turned on only in 3 out of the 12 datasets (Yan, Biase and Ting), because the remaining 7 datasets would each have zero genes surviving its quality filtering. For SC3 and t-SNE + *k*-means, all reported results are from random seed 123.

**Table 1.**
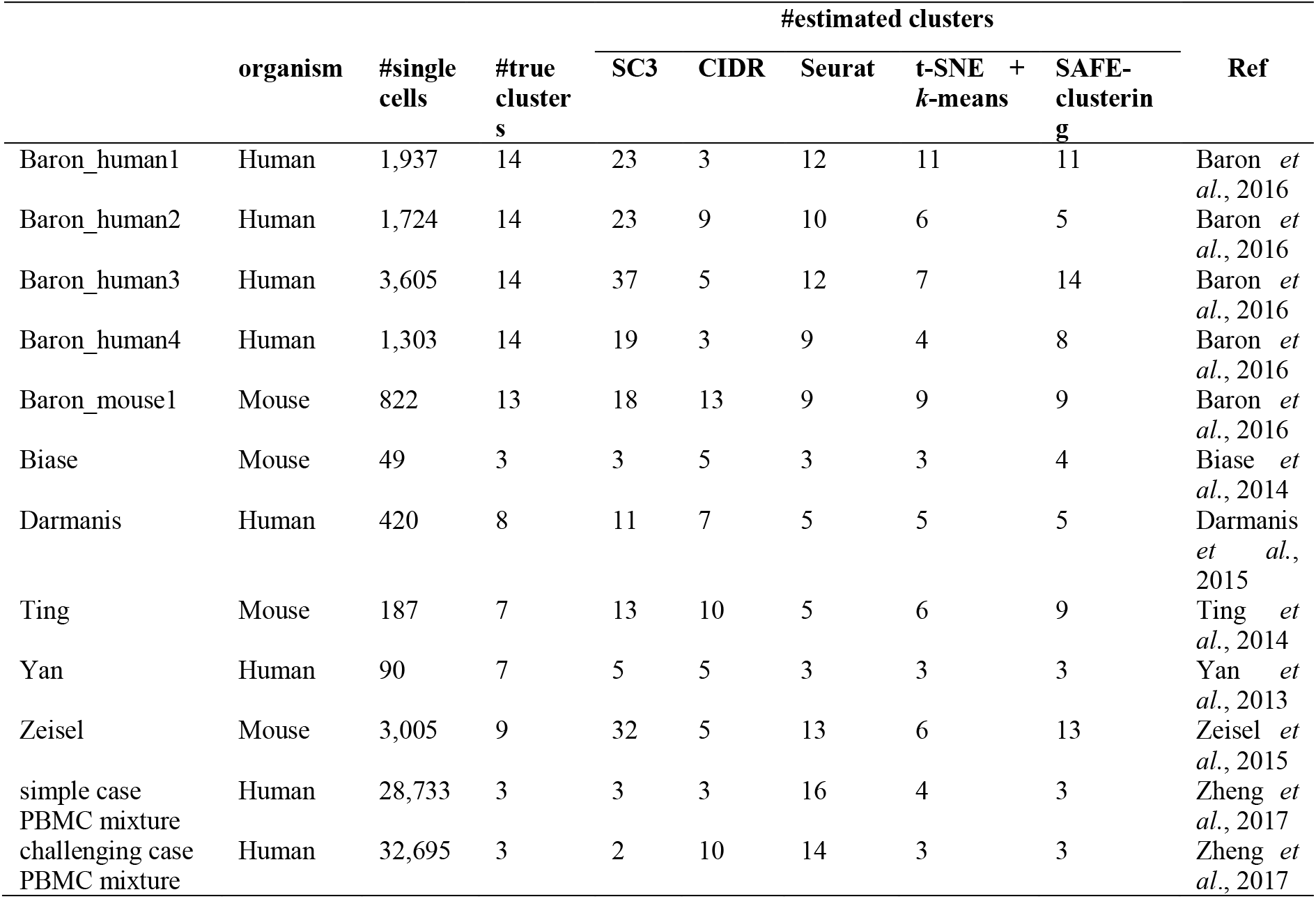
Major characteristics of the 12 benchmarking datasets, including organism origin, number of single cells, the numbers of true and estimated clusters by SAFE-clustering and four individual methods, as well as references.

Performance is measured by the similarity between the estimated cluster labels (*L_E_*) and the true cluster labels (*L_T_*) using the Adjusted Rand Index (ARI) (Hubert and Arabie, 1985):

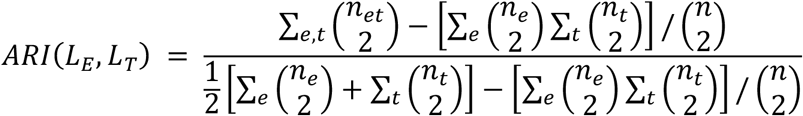

where *n* is the total number of single cells; *n_e_* and *n_t_* are the number of single cells in estimated cluster *e* and in true cluster *t*, respectively; and *n_et_* is the number of single cells shared by estimated cluster *e* and true cluster *t*. ARI ranges from 0 to 1, where 1 means the estimated cluster is exactly same to the true cluster, while 0 means the two are completely different.

Computing time reported in this work is all from running on an iMac with 3.4 GHz Intel Core 1.5, 32 GB 1600 MHz DDR3 of RAM and OS X 10.9.5 operating system.

## 3 RESULTS

### 3.1 Individual methods capture different characteristics of scRNA-Seq data

We observe relatively moderate similarity among solutions from individual ensemble methods (Fig. 2), consistent with findings from Freytag *et al*. (2017). These may reflect different methods capturing different aspects of information from the rather complex and highdimensional scRNA-Seq data, leading to different solutions, but no clear winner.

**Fig. 2.**
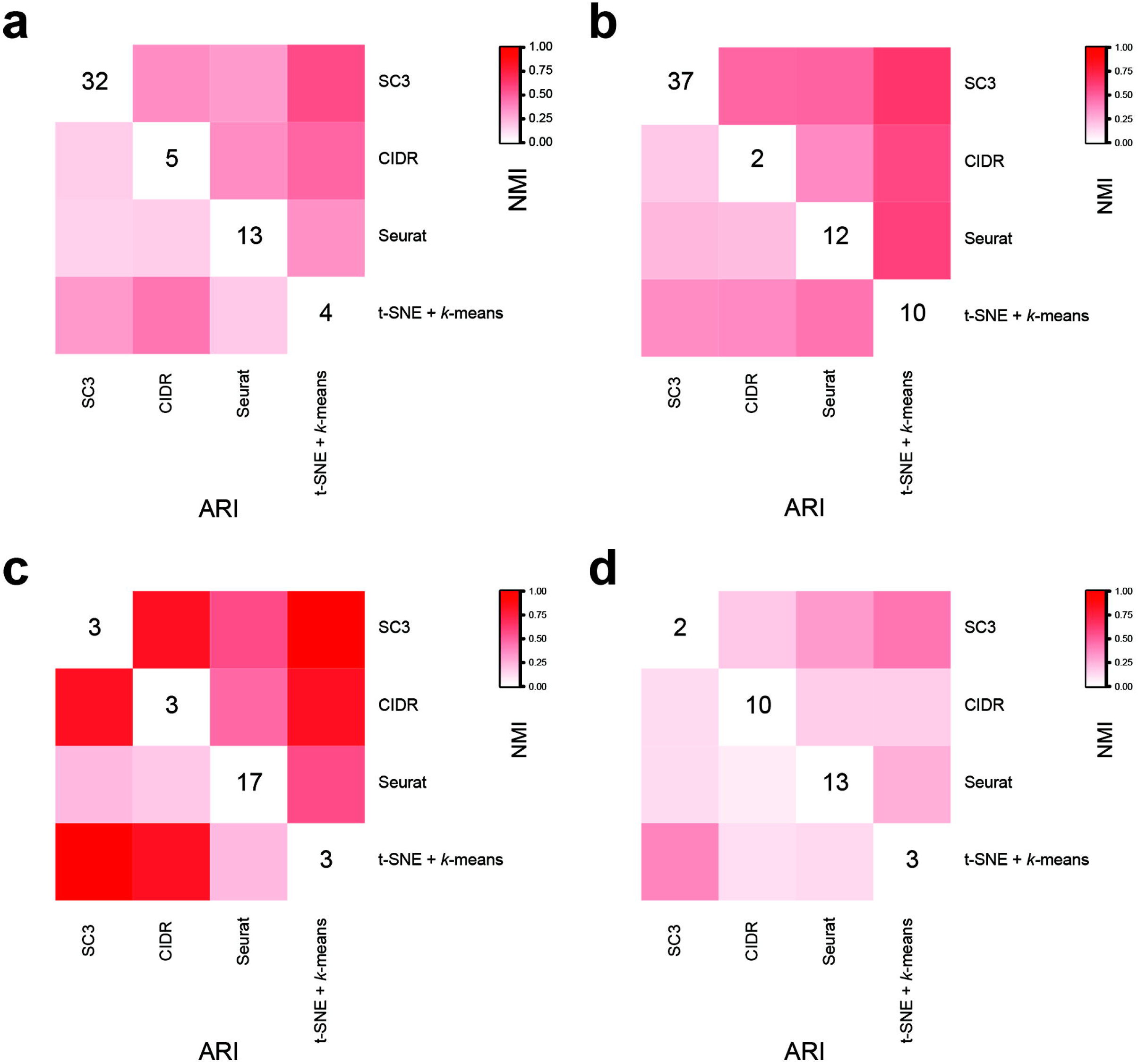
Similarity of solutions from individual clustering methods. (**a**) Zeisel dataset; (**b**) Baron_human3 dataset; (**c**) simple case PBMC mixture dataset; (**d**) challenging case PBMC mixture dataset.

### 3.2 Improving and running individual ensemble methods

#### Seurat

Seurat provides a “resolution” parameter to alter the granularity of the clustering results. However, the default “resolution” (= 0.8) tends to result in no clustering for small datasets, as shown in the SC3 paper (Kiselev *et al*., 2017). To further evaluate the performance of Seurat on small datasets, we generated 100 subsets of samples from the Darmanis dataset, using stratified random sampling without replacement where each cell type was one stratum and single cells from each cell type were randomly selected according to the corresponding cell type proportion. Our sampling strategy resulted in 61 - 239 single cells from the eight cell types, across the 100 generated datasets. The resolution was set to 0.6, 0.9 and 1.2, respectively, following the instruction of Seurat. Due to non-determination from random sampling, the sampling process and the downstream clustering were repeated 100 times for each resolution. The performance of different resolution is quantified by ARI according to published clustering. When sample size ranges from 61 to 150, Seurat clustering with resolution = 1.2 performs significantly better than 0.6 and 0.9 (*p* < 0.05, **Supplementary Fig. S1a**), except for the case between resolution 0.9 and 1.2 in the subset of 120 cells (*p* = 0.124). Comparatively, only one cluster is identified in the subset of 61 cells when resolution = 0.6. When sample size increases to 210, resolution makes no difference.

When applying Seurat to the three small datasets, Biase (*n* = 49 single cells), Yan (*n* = 90) and Ting (*n* = 187), we used all three resolutions. Overall, Seurat performed better with resolution =1.2 (**Supplementary Fig. S1b**), with the exception of Yan dataset, where clusterings with all the three resolutions are the same. For Biase dataset, Seurat cannot distinguish different cell types with resolution = 0.6, but ARI reaching to 1 when resolution increases to 1.2.

#### tSNE + *k*-means

Results from t-SNE + *k*-means are stochastic rather than deterministic. To mitigate the fluctuations across runs, we used the ADPclust R-package (Wang and Xu, 2015) to first obtain clustering centroids. We compared the performance with and without this ADPclust centroid estimation step before *k*-means, on four datasets, Yan, Ting, Darmanis and Baron_human2. Expression matrix was log-transformed and dimensionality reduced using t-SNE. For each clustering strategy, t-SNE was carried out 100 times. The number of clusters ranged from 2 to (*k_M_* + 2), where *k_M_* is the maximum number of clusters, in term of the true and estimated numbers of clusters. As expected, ARI’s from the 100 datasets without ADPclust centroid estimation varied dramatically at most *k*’s attempted where *k* is the number of clusters fed to *k*-means (**Supplementary Fig. S2**). In contrast, with ADPclust centroid the estimation had much improved stability.

#### SC3

For the two PBMC mixture datasets, SC3 estimated 588 and 586 clusters for the simple and challenging case, respectively, dramatically deviating from the truth (*k* = 3 for both two datasets). The *k* estimation method in SC3 has not been benchmarked and validated for large, shallowly sequenced datasets, and it is likely that the distribution of eigenvalues of the covariance matrix does not adhere to the assumed Tracy-Widom distribution (Tracy and Widom, 1994). However, clustering results of SC3 are not affected by this since *k* estimation in SC3 is completely independent of the clustering algorithm (SC3 source codes on Dec 11, 2017; https://github.com/hemberg-lab/SC3/blob/8478ff2c8f523f004d129aec56ae57ce6853bd12/R/CoreFunctions.R). We therefore performed PCA plot visualization (using *plotPCA* function of scater R-package) to narrow down a reasonable range of *k*. PCA plot suggested 3 distinct clusters for the simple case and 2 clusters for the challenging case (**Supplementary Fig. S3**). We therefore decided, for SC3, on *k* = 3 for the simple case and *k* = 2 for the challenging case. SC3 ARI for the simple case at our selected *k* = 3 is 0.995 and for the challenging case at *k* = 2 is 0.595.

Because of the issue revealed from the PBMC mixture datasets and because estimation of number of clusters can be separated from clustering *per se*, we ran SC3 for both datasets within a more reasonable range of *k*: from 2 to 7. Using the SC3 results from this range, we assessed the robustness of our SAFE-clustering method, holding all the other three individual methods constant. **Supplementary Fig. S4** shows that ARI from SC3 fluctuates considerably (0.599 - 0.995 and 0.596 - 0.768 for the simple and challenging case, respectively) when *k* increases from 2 to 7. Comparatively, results from our SAFE-clustering are much more stable (ARI ranges from 0.852 to 0.995 for the simple case and from 0.582 to 0.694 for the challenging case, respectively). These results suggest that even with a non-optimal *k* selected by one individual method, our SAFE-clustering ensemble method is able to generate robustly accurate results, because our ensemble method borrows information from the other contributing methods. Furthermore, SAFE-clustering correctly estimates the number of clusters (i.e., 3) for both the simple and the challenging case with SC3’s *k* ranging from 2 to 7.

### 3.3 Benchmarking of SAFE-clustering across 12 datasets

We benchmarked SAFE-clustering together with its four embedded individual clustering methods on 12 published scRNA-Seq datasets, reflecting a wide spectrum of experimental technology, sequencing depth, tissue origin, number and heterogeneity of single cells examined (details are summarized in Table 1 and **Supplementary Table 1**). Among the 12 datasets, we examine two large peripheral blood mononuclear cells (PBMC) mixture datasets with >28,000 single cells were constructed by mixing single-cell datasets of purified cell types generated by the 10× Genomics (Zheng *et al*., 2017) as described in Sun *et al*. (2017). Specifically, we created one dataset representing a “simple case” with 28,733 single cells from three distinct cell types: CD56+ natural killer cells, CD19+ B cells and CD4+/CD25+ regulatory T cells; and the other dataset representing a “challenging case” with 32,695 single cells from three highly similar cell types: CD8+/CD45RA+ naive cytotoxic T cells, CD4+/CD45RA+/CD25-naive T cells and CD4+/CD25+ regulatory T cells.

For the 12 datasets attempted, SAFE-clustering outperforms all the individual solutions in five datasets: Baron_human1, Baron_human3, Baron_mouse1, and the two PBMC mixture datasets (Fig. 3). Furthermore, SAFE-clustering performs better than at least two individual methods in six additional datasets (Biase, Yan, Darmanis, Zeisel, and Baron_human2 and 4) (Fig. 3). These results show that SAFE-clustering performs robustly well across various datasets. We also compared the estimated number of clusters and found that among individual methods, CIDR performs the best (Fig. 4b); SC3 tends to overestimate the number of clusters (Fig. 4a), while t-SNE + *k*-means tends to underestimate (Fig. 4d). Our SAFE-clustering outperforms all individual solutions (Fig. 4e and f), quantified by the average absolution deviation from the true/gold-standard cluster numbers (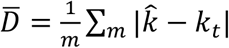, where *m* is the number of datasets (= 12 in our work); 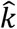 is the estimated number of clusters; and *k_t_* is the true (or predefined gold/silver standard) number of cell types.

**Fig. 3.**
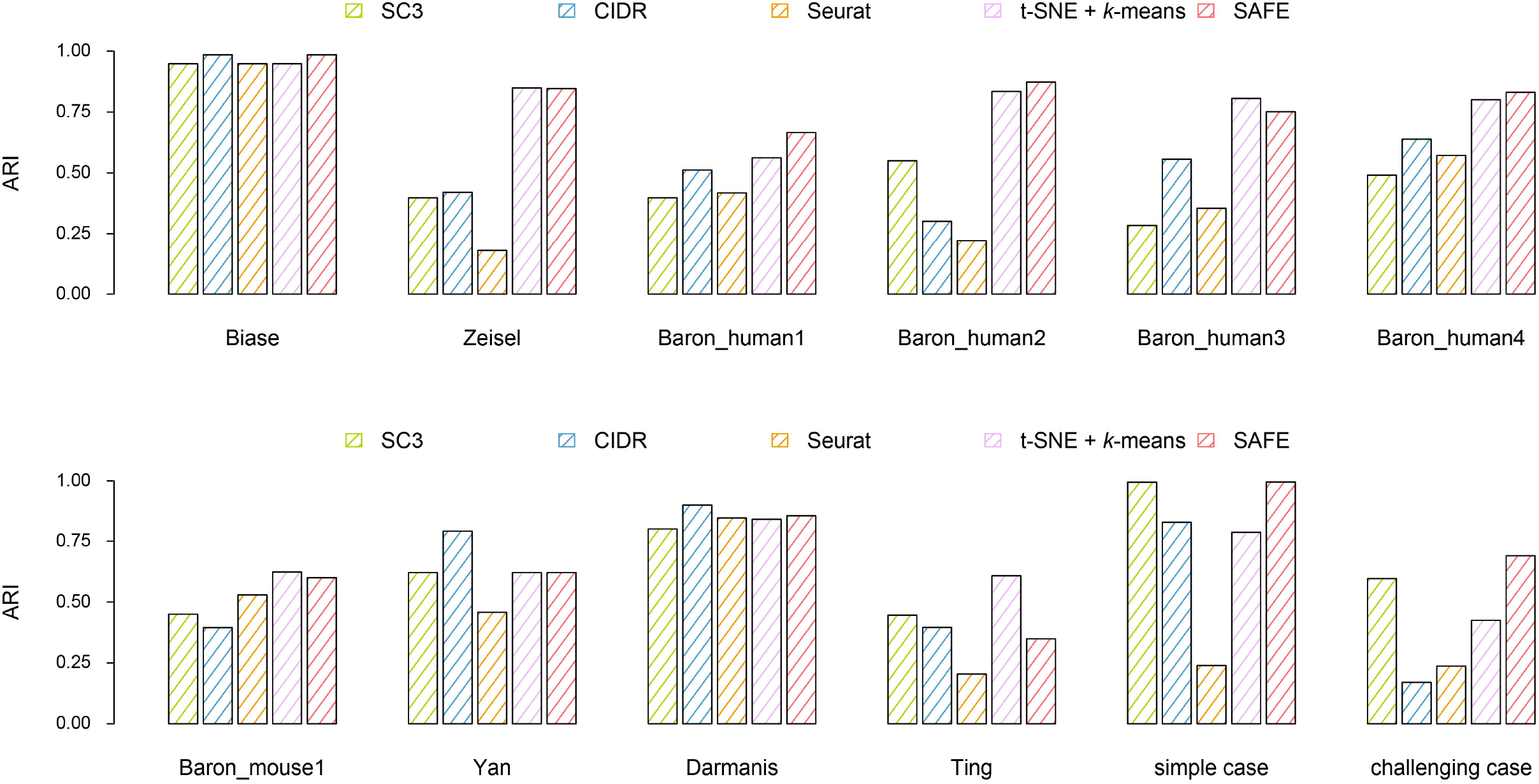
Benchmarking of SAFE-clustering in 12 published datasets. Adjusted Rand Index (ARI) is employed to measure the similarity between inferred and true cluster labels. Detailed information of the 12 datasets can be found in Table 1 and **Supplementary Table 1**.

**Fig. 4.**
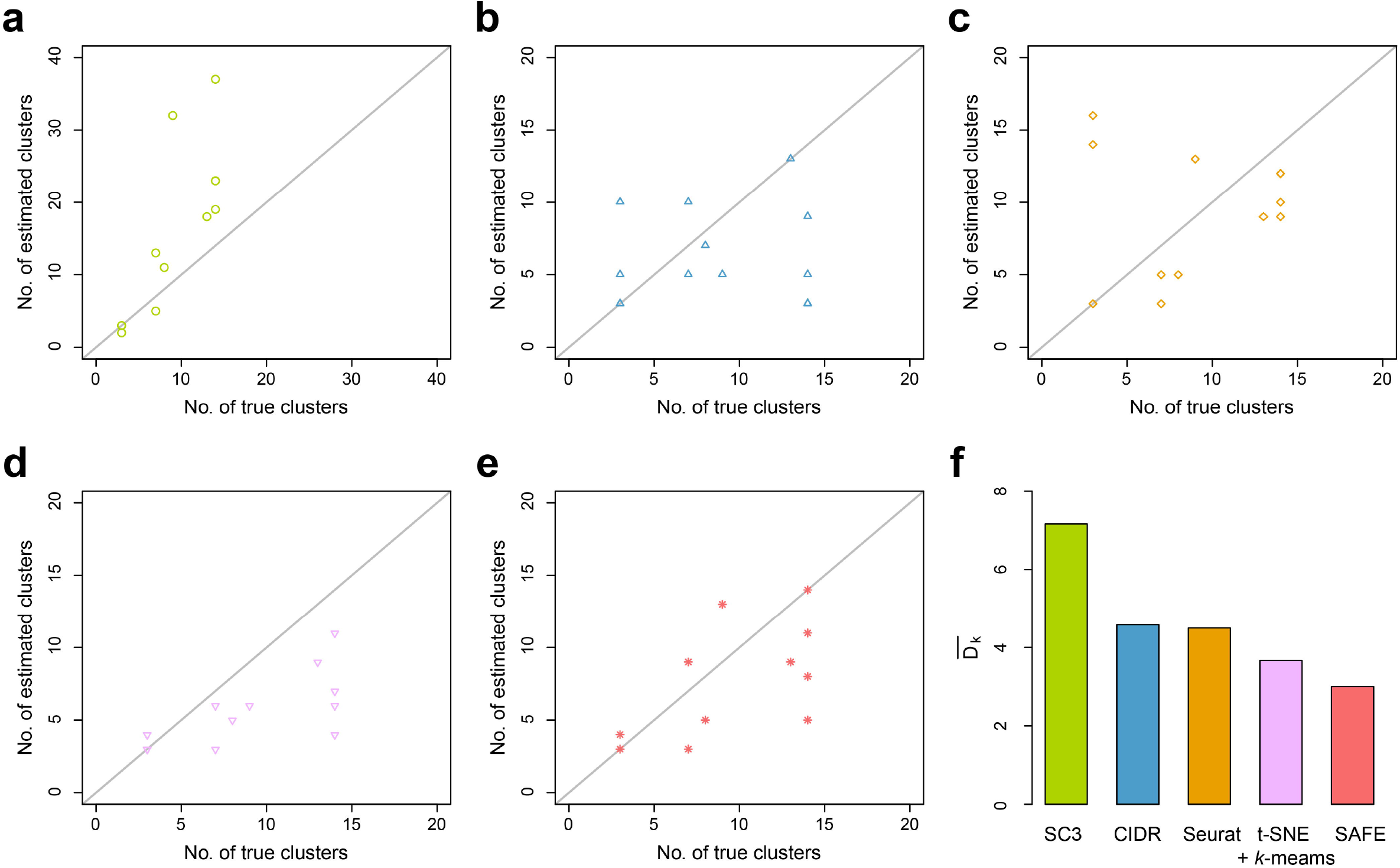
Accuracy evaluation of the inferred number of clusters. (**a-e**) Correlations between inferred cluster numbers from SC3, CIDR, Seurat, t-SNE + *k*-means and SAFE-clustering, respectively, and the true cluster numbers, across the 12 benchmarking datasets. (**f**) Average deviations between the inferred and the true numbers of clusters, measured by 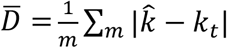, where the number of datasets *m* equals to 12.

For the simple case PBMC mixture dataset, both CIDR and SC3 yielded 3 clusters with Adjusted Rand Index (ARI) of 0.827 and 0.995, respectively (Fig. 3). Seurat assigned the single cells into 16 clusters with an ARI of 0.239. Also, Seurat failed to generate clustering results for three (out of 28,733) single cells because of <200 expressed genes in these cells. For t-SNE + *k*-means, we applied t-SNE on the top 1,000 most variable genes to save computing time and memory usage (**Supplementary Fig. 5**), identifying three clusters with an ARI of 0.976. Combining the four individual solutions, SAFE-clustering generated the most accurate result with an ARI of 0.995 (Fig. 3 and **Supplementary Fig. 6a**). Moreover, all the three single cells not clustered by Seurat were correctly assigned into their corresponding clusters by SAFE-clustering’s borrowing information from the remaining three individual solutions.

For the challenging case PBMC mixture dataset, none of the four individual methods performed well, because CD4+/CD45RA+/CD25-naive T cells are quite similar to CD4+/CD25+ regulatory T cells. SC3 generated the most accurate individual solution, identifying two clusters with an ARI of 0.595 (Fig. 3), followed by t-SNE + *k*-means (3 clusters and ARI = 0.405). Similar to the simple case, Seurat failed to generate clustering results for 28 single cells with <200 expressed genes, and resulted in 13 clusters with an ARI of 0.264. SAFE-clustering again outperformed all the four individual methods (Fig. 3 and **Supplementary Fig. 6b**), correctly identifying three clusters with an ARI = 0.612, and correctly clustering 23 out of the 28 single cells which were not clustered by Seurat. These results strongly suggest that SAFE-clustering can provide robust and high-quality clustering even under challenging scenarios.

Besides the four individual methods used in our package, we also compared our results with two additional widely-used clustering methods SIMLR (Wang *et al*., 2017) and RaceID (Grün *et al*., 2015), and the results showed that SAFE-clustering excels SIMLR in 8 out of 12 datasets, and outperforms than RaceID in 11 out of 12 datasets (**Supplementary Fig. S7**). To assess the extensibility of SAFE-clustering to other scRNA-seq clustering methods, we incorporated one more individual method, SIMLR, into our SAFE-clustering and found that the ensemble solutions are similar to those from the original SAFE-clustering without the fifth SIMLR method (**Supplementary Fig. S7**). Our results suggest SAFE-clustering is robust also to the increasing number of employed individual methods.

Additionally, we evaluated the potential impacts of several factors: inclusion/exclusion of ribosomal protein genes, filtering on percentage of mitochondrial reads, dropout imputation and denoising of expression profiles, perplexity parameter for t-SNE, and number of t-SNE dimensions carried forward for *k*-means clustering. Details of the evaluation results are given in the **Supplementary Results**.

Overall, across the 12 datasets, SAFE-clustering on average improved ARI by 36.0% over the average of the individual methods, and up to 18.5% over the best individual method for each dataset. All codes for two example datasets are made available via an R markdown freely available at both https://github.com/yycunc/SAFEclustering and https://yunliweb.its.unc.edu/safe/SAFEclustering_tutorial.html.

### 3.4 Benchmarking of three hypergraph partitioning algorithms in SAFE-clustering

SAFE-clustering has three hypergraph partitioning algorithms implemented. Among them, CSPA is computationally expensive for datasets with large number of single cells because computational complexity increases quadratically with the number of single cells (Punera and Ghosh, 2008). To assess the feasibility of the three algorithms on big datasets, we recorded the running time for the simple case of 28,733 cells. As the running time is insensitive to the number of clusters *k*, a 3-way partitioning (that is, *k* was set at 3, the true cluster number) was performed, running each of the algorithms 100 times. As expected, HGPA is ultrafast taking an average of 0.51 +/− 0.02 *second per clustering* (*s/c*), followed by MCLA, 8.26 +/− 1.54 *s/c*. CSPA is the slowest with ~576.64 +/− 0.74 *s/c* (Fig. 5a). Finally and importantly, we would like to note that computational costs of these ensemble algorithms are negligible (HGPA and MCLA) or low (CSPA), compared to the computing costs of individual clustering methods (2.5 - 22 hours per clustering).

**Fig. 5.**
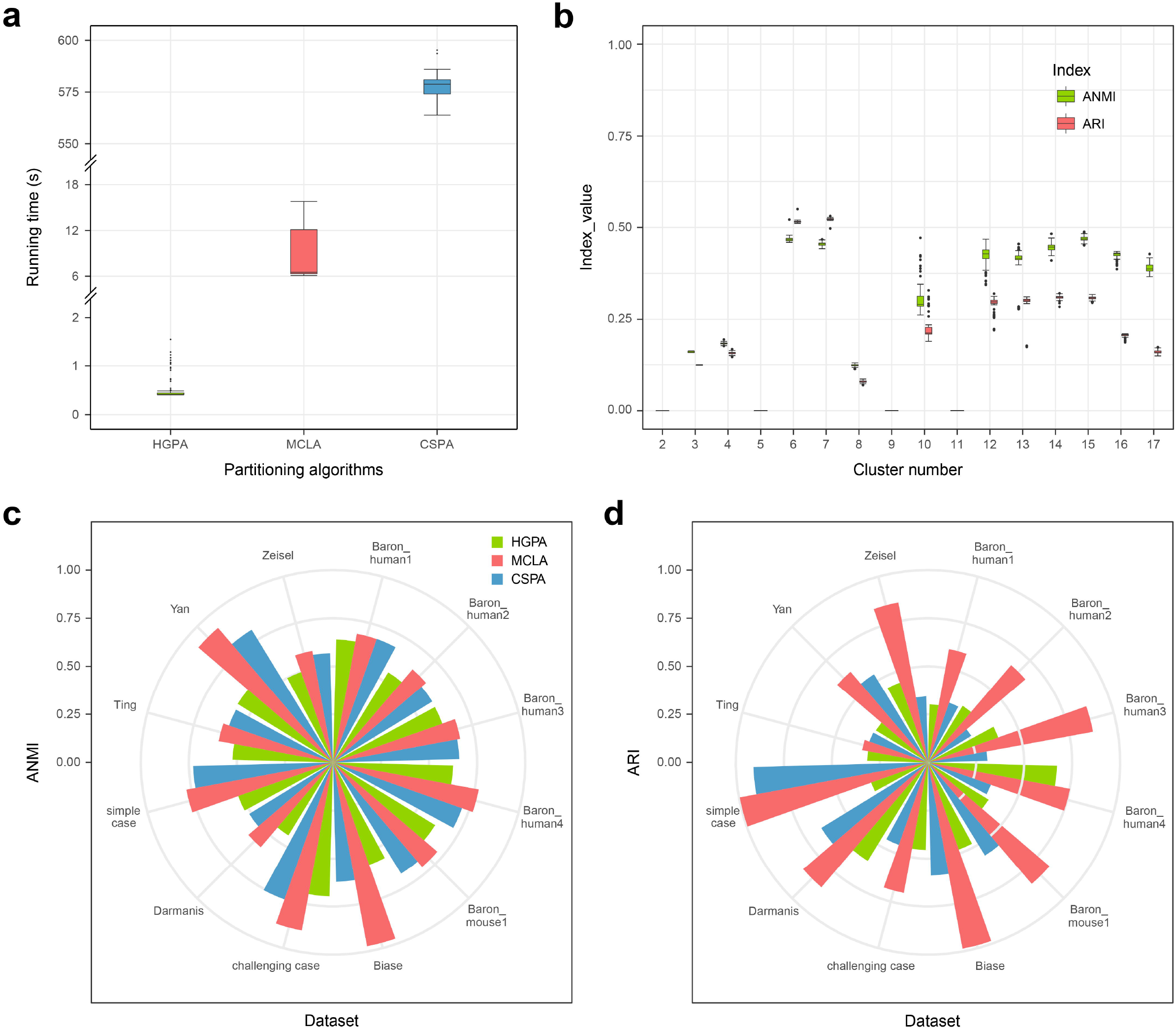
Benchmark of the three hypergraph partitioning algorithms: HGPA, MCLA and CSPA. (**a**) Running time for 3-way partitioning of simple case PBMC mixture dataset with 28,733 single cells using each of the three partitioning algorithms. Each algorithm was applied 100 times. (**b**) Stability of HGPA from 100 runs using simple case PBMC mixture dataset with 28,733 single cells. (**c**) Similarity between consensus clustering and individual solutions in 12 benchmarking datasets, measured by Average Normalized Mutual Information (ANMI). (**d**) Performance of the three partitioning algorithms, measured by ARI, across the 12 benchmarking datasets.

Among the three ensemble algorithms, MCLA and CSPA results are deterministic conditional on any specified random number generator (RNG) seed. HGPA, however, generates stochastic results even with a specified RNG seed. To evaluate the stability of HGPA’s clustering results, we performed HGPA partitioning 100 times on the simple case dataset and calculated both ANMI and ARI for each run. Fig. 5b shows that HGPA results, although relatively stable, vary slightly across different runs. Another consequence of HGPA’s stochasticity is that different numbers of cluster may be estimated. Therefore, SAFE-clustering by default runs HGPA 10 times, selects the run with the median ANMI value among the 10 runs, and outputs the corresponding consensus result.

To evaluate the performance of the three hypergraph partitioning algorithms, we performed ensemble clustering of the 12 datasets using each of them (namely HGPA, MCLA and CSPA) separately. Comparatively, MCLA is a clear winner: manifesting the highest ANMI in 11 out of the 12 benchmarking datasets (Fig. 5c); and exhibiting the highest ARI in 11 out of the 12 datasets (Fig. 5d). For the single dataset (Baron_human3) where MCLA is not the best according to ANMI, its ANMI (0.658) is a close match of the best (0.662 from CSPA). In addition, in this Baron_human3 dataset, if gauged using ARI, MCLA again outperforms all other methods with ARI = 0.507 and the second best ARI = 0.215 from CSPA. For the Ting dataset where MCLA is not the best according to the ARI metric, it is the close match second best with ARI = 0.429, compared with the best (from CSPA) with ARI = 0.556 and 0.465 respectively. These results suggest that MCLA provides more accurate consensus clustering than the other two algorithms. Therefore, SAFE-clustering uses MCLA as the default partitioning algorithm. These three partitioning algorithms vary in performance due to their inherently differences: although they all employ hyperedges and hypergraphs, they differ quite drastically in *how* (Karypis and Kumar, 1998; Karypis *et al*., 1999). Specifically, HGPA partitions the hypergraph by cutting a minimal number of hyperedges that creates k clusters of approximately equal size, which would not be optimal when cluster sizes vary substantially. CSPA starts with a similarity matrix computed from the hypergraph to perform partitioning, and MCLA first computes a pairwise Jaccard similarity matrix and collapses related hyperedges(clusters).

## 4 DISCUSSION

We present SAFE-clustering, an unsupervised ensemble method to provide fast, accurate and flexible clustering for scRNA-Seq data. Although there are a number of clustering methods developed for scRNA-Seq data in the recent literature, individual clustering methods differ in many aspects including data pre-processing, choice of distance metrics, clustering method, and model selection to determine number of clusters, thus their performances tend to vary, sometimes rather dramatically, across datasets. There is no clear winner among the existing clustering methods. Our SAFE-clustering employs hypergraph portioning algorithms to build an ensemble solution based on multiple solutions from individual clustering methods, the first time ensemble has been leveraged across different types of methods for scRNA-Seq data. The leveraged information from all these individual methods enables our SAFE Ensemble method to reach robustly satisfactory performance across datasets. We have benchmarked SAFE-clustering along with four individual clustering methods (SC3, CIDR, Seurat and t-SNE + *k*-means) on 12 published scRNA-Seq datasets, which is the most comprehensive to date. Among the 12 datasets, SAFE-clustering outperforms all four individual solutions in five benchmarking datasets, and performs better than at least two individual methods in six datasets (Fig. 3). For the two PBMC mixture datasets with 28,733 and 32,695 single cells respectively, SAFE-clustering accurately identifies the three cell types of ARI = 0.995 and 0.612 respectively (Fig. 3 and **Supplementary Fig. S6**). Gauged by ARI, SAFE-clustering outperforms the most accurate existing method for each dataset by up to 18.5%, and on average by 36.0% over the average performance of the four state-of-the-art methods, across the 12 datasets. Moreover, although care needs to be taken for interpreting these cluster number estimates (**Supplementary results** and **Supplementary Fig. S8**), SAFE-clustering provides the most accurate estimation on the number of cell types compared to the individual methods: SAFE-clustering’s average absolute deviation from true cluster numbers (3.58) is substantially smaller than that any of the four individual methods (average absolute deviation ranging from 4.42 to 7.17) (Fig. 4f). These results suggest that SAFE-clustering produces more stable and accurate clustering across various datasets. We note that many pre-processing steps can also influence results noticeably and should be carried out with caution. We have made efforts to evaluate a number of them (**Supplementary results**) and add corresponding options to our SAFE-clustering package with default values. A complete evaluation of all possible preprocessing choices is beyond the scope of this work, if not impossible. Finally, SAFE-clustering is computationally efficient, with the additional hypergraph partitioning of individual methods’ cluster assignments taking less than 10 seconds to cluster 28,733 cells, using the default MCLA algorithm (Fig. 5a). SAFE-clustering is scalable to even larger datasets; taking 5-22 minutes for datasets with 150,000 - 300,000 single cell for instance (**Supplementary Fig. S9**). We anticipate that SAFE-clustering will prove valuable for increasingly larger number of investigators working with scRNA-Seq data.

## ACKNOWLEDGEMENT

The authors would like to thank Drs. Ming Hu and Sullivan for helpful discussions on the software and on earlier versions of the manuscript. We also want to acknowledge the various studies from which the benchmarking datasets were generated. We would also like to thank all Li Lab members for providing feedback and suggestions on the work while in progress; and for testing the software towards the end of the project.

## REFERENCES

Arsenio, J. et al. (2014) Early specification of CD8+ T lymphocyte fates during adaptive immunity revealed by single-cell gene-expression analyses. Nat. Immunol., 15, 365–372.

Baron, M. et al. (2016) A single-cell transcriptomic map of the human and mouse pancreas reveals inter-and intra-cell population structure. Cell Syst., 3, 346–360.

Biase, F.H. et al. (2014) Cell fate inclination within 2-cell and 4-cell mouse embryos revealed by single-cell RNA sequencing. Genome Res., 24, 1787–1796.

Buganim, Y. et al. (2012) Single-cell expression analyses during cellular reprogramming reveal an early stochastic and a late hierarchic phase. Cell, 150, 1209–1222.

Caliński, T. and Harabasz, J. (1974) A dendrite method for cluster analysis. Commun. Stat. Methods, 3, 1–27.

Darmanis, S. et al. (2015) A survey of human brain transcriptome diversity at the single cell level. Proc. Natl. Acad. Sci., 112, 7285–7290.

Freytag, S. et al. (2017) Cluster Headache: Comparing Clustering Tools for 10X Single Cell Sequencing Data. bioRxiv, 203752.

Ghosh, J. and Acharya, A. (2011) Cluster ensembles. Wiley Interdiscip. Rev. Data Min. Knowl. Discov., 1, 305–315.

Grün, D. et al. (2015) Single-cell messenger RNA sequencing reveals rare intestinal cell types. Nature, 525, 251–255.

Hubert, L. and Arabie, P. (1985) Comparing partitions. J. Classif., 2, 193–218.

Hubner, I.A. et al. (2005) High-resolution protein folding with a transferable potential. Proc. Natl. Acad. Sci. U. S. A., 102, 18914–18919.

Jaitin, D.A. et al. (2014) Massively parallel single-cell RNA-seq for marker-free decomposition of tissues into cell types. Science (80-.)., 343, 776–779.

Jia, C. et al. (2017) Accounting for technical noise in differential expression analysis of single-cell RNA sequencing data. Nucleic Acids Res., 45, 10978–10988.

Kalisky, T. and Quake, S.R. (2011) Single-cell genomics. Nat. Methods, 8, 311–314.

Karypis, G. et al. (1999) Multilevel hypergraph partitioning: applications in VLSI domain. IEEE Trans. Very Large Scale Integr. Syst., 7, 69–79.

Karypis, G. and Kumar, V. (1998) A fast and high quality multilevel scheme for partitioning irregular graphs. SIAM J. Sci. Comput., 20, 359–392.

Kiselev, V.Y. et al. (2017) SC3: consensus clustering of single-cell RNA-seq data. Nat. Methods, 14, 483–486.

Kuepfer, L. et al. (2007) Ensemble modeling for analysis of cell signaling dynamics. Nat. Biotechnol., 25, 1001–1006.

Lin, P. et al. (2017) CIDR: Ultrafast and accurate clustering through imputation for single-cell RNA-seq data. Genome Biol., 18, 59.

Van der Maaten, L. and Hinton, G. (2008) Visualizing data using t-sne. J. Mach. Learn. Res., 9, 2579–2605.

Macosko, E.Z. et al. (2015) Highly parallel genome-wide expression profiling of individual cells using nanoliter droplets. Cell, 161, 1202–1214.

Mahata, B. et al. (2014) Single-cell RNA sequencing reveals T helper cells synthesizing steroids de novo to contribute to immune homeostasis. Cell Rep., 7, 1130–1142.

Patterson, N. et al. (2006) Population structure and eigenanalysis. PLoS Genet., 2, e190.

Punera, K. and Ghosh, J. (2008) Consensus-based ensembles of soft clusterings. Appl. Artif. Intell., 22, 780–810.

Rozenblatt-Rosen, O. et al. (2017) The Human Cell Atlas: from vision to reality. Nature.

Satija, R. et al. (2015) Spatial reconstruction of single-cell gene expression data. Nat. Biotechnol., 33, 495–502.

Shalek, A.K. et al. (2013) Single-cell transcriptomics reveals bimodality in expression and splicing in immune cells. Nature, 498, 236–240.

Shin, J. et al. (2015) Single-cell RNA-seq with waterfall reveals molecular cascades underlying adult neurogenesis. Cell Stem Cell, 17, 360–372.

Strehl, A. and Ghosh, J. (2002) Cluster ensembles---a knowledge reuse framework for combining multiple partitions. J. Mach. Learn. Res., 3, 583–617.

Sun, Z. et al. (2017) DIMM-SC: A Dirichlet mixture model for clustering droplet-based single cell transcriptomic data. Bioinformatics, btx490.

Tang, F. et al. (2010) Tracing the derivation of embryonic stem cells from the inner cell mass by single-cell RNA-Seq analysis. Cell Stem Cell, 6, 468–478.

Ting, D.T. et al. (2014) Single-cell RNA sequencing identifies extracellular matrix gene expression by pancreatic circulating tumor cells. Cell Rep., 8, 1905–1918.

Tracy, C.A. and Widom, H. (1994) Level-spacing distributions and the Airy kernel. Commun. Math. Phys., 159, 151–174.

Trapnell, C. et al. (2014) The dynamics and regulators of cell fate decisions are revealed by pseudotemporal ordering of single cells. Nat. Biotechnol., 32, 381–386.

Treutlein, B. et al. (2014) Reconstructing lineage hierarchies of the distal lung epithelium using single-cell RNA-seq. Nature, 509, 371–375.

Waltman, L. and van Eck, N.J. (2013) A smart local moving algorithm for large-scale modularity-based community detection. Eur. Phys. J. B, 86, 471.

Wang, B. et al. (2017) Visualization and analysis of single-cell RNA-seq data by kernel-based similarity learning. Nat. Methods, 14, 414–416.

Wang, X.-F. and Xu, Y. (2015) Fast clustering using adaptive density peak detection. Stat. Methods Med. Res., 962280215609948.

Wolf, F.A. et al. (2018) SCANPY: large-scale single-cell gene expression data analysis. Genome Biol., 19, 15.

Yan, L. et al. (2013) Single-cell RNA-Seq profiling of human preimplantation embryos and embryonic stem cells. Nat. Struct. Mol. Biol., 20, 1131–1139.

Zeisel, A. et al. (2015) Cell types in the mouse cortex and hippocampus revealed by single-cell RNA-seq. Science (80-.)., 347, 1138–1142.

Zhang, J.M. et al. (2018) An interpretable framework for clustering single-cell RNA-Seq datasets. BMC Bioinformatics, 19, 93.

Zheng, G.X.Y. et al. (2017) Massively parallel digital transcriptional profiling of single cells. Nat. Commun., 8, 14049.

Zhu, L. et al. (2017) A Unified Statistical Framework for Single Cell and Bulk RNA Sequencing Data. bioRxiv, 206532.

