## Supplementary Materials for "SAFE-clustering: Single-cell Aggregated (From Ensemble) Clustering for Single-cell RNA-seq Data" (from another version of this preprint: version 2, 2018-03-28)

**Supplementary File for Online Publication**

**Supplementary Table S1.** Major characteristics of the 14 benchmarking datasets. Table 1 lists several major characteristics of the 14 benchmarking datasets, including tissue origin, technology, datatype, and sequencing depth.

| **Dataset** | **Tissue origin** | **Technology** | **Datatype** | **Median/mean (range) sequencing per single cell*** |
| --- | --- | --- | --- | --- |
| Baron-human1 | Human pancreas | inDrop | UMI | 5,346/5,805 (1.3-32K) unique transcripts |
| Baron-human2 | Human pancreas | inDrop | UMI | 4,890/5,109 (1.3-15K) unique transcripts |
| Baron-human3 | Human pancreas | inDrop | UMI | 4,742/5,859 (1.2-30K) unique transcripts |
| Baron-human4 | Human pancreas | inDrop | UMI | 6,017/6,729 (1.4-35K) unique transcripts |
| Baron-mouse1 | Mouse pancreas | inDrop | UMI | 2,937/3741 (1-16K) unique transcripts |
| Biase | Mouse embryo | SMART-Seq SMARTer | FPKM | Average of 37.9 million 100bp reads^**^ |
| Darmanis | Human brain | SMART-Seq SMARTer | Gene counts | Average of 2.8 million reads^**^ 0.5/0.6 (0.1-2.0) million reads |
| Ting | Mouse tumors | scRNA-Seq (modified Tang2010 protocol^1^) | CPM | Average 4.4-8.5 million reads^**^ |
| Yan | Human embryo | scRNA-Seq (Tang2010 protocol^1^) | RPKM | Average of 35.3 million 100bp reads^**^  26.5/28.3 (14.2-50.2) million reads |
| Zeisel | Mouse brain | Quantitative scRNA-seq with UMI^2^ | UMI | 12.1K/14.0K (2.5-62.0K) UMIs |
| Challenging case | PBMC | 10X Genomics GemCode | UMI | 1,288/1,357 (0.3-7.3K) unique transcripts |
| Simple case | PBMC | 10X Genomics GemCode | UMI | 1,337/1,459 (0.4-7.3K) unique transcripts |

^*^: Re-calculated using the data we obtained if not otherwise specified

^**^: Based on original publication (references provided in Table 1)

1. Tang, F., Barbacioru, C., Nordman, E., Li, B., Xu, N., Bashkirov, V. I., . . . Surani, M. A. (2010). RNA-Seq analysis to capture the transcriptome landscape of a single cell. Nature protocols, 5(3), 516-535.
2. Islam, S., Zeisel, A., Joost, S., La Manno, G., Zajac, P., Kasper, M., . . . Linnarsson, S. (2014). Quantitative single-cell RNA-seq with unique molecular identifiers. Nature methods, 11(2), 163-166.

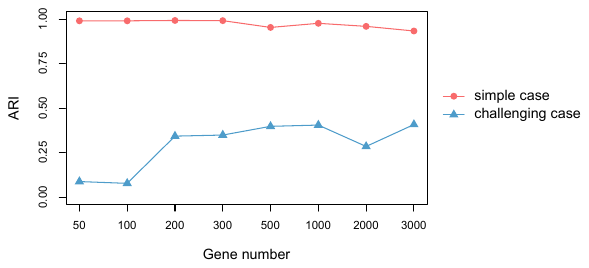

**Supplementary Fig. S1.** ARI of tsne+kmeans including the top most variable genes.

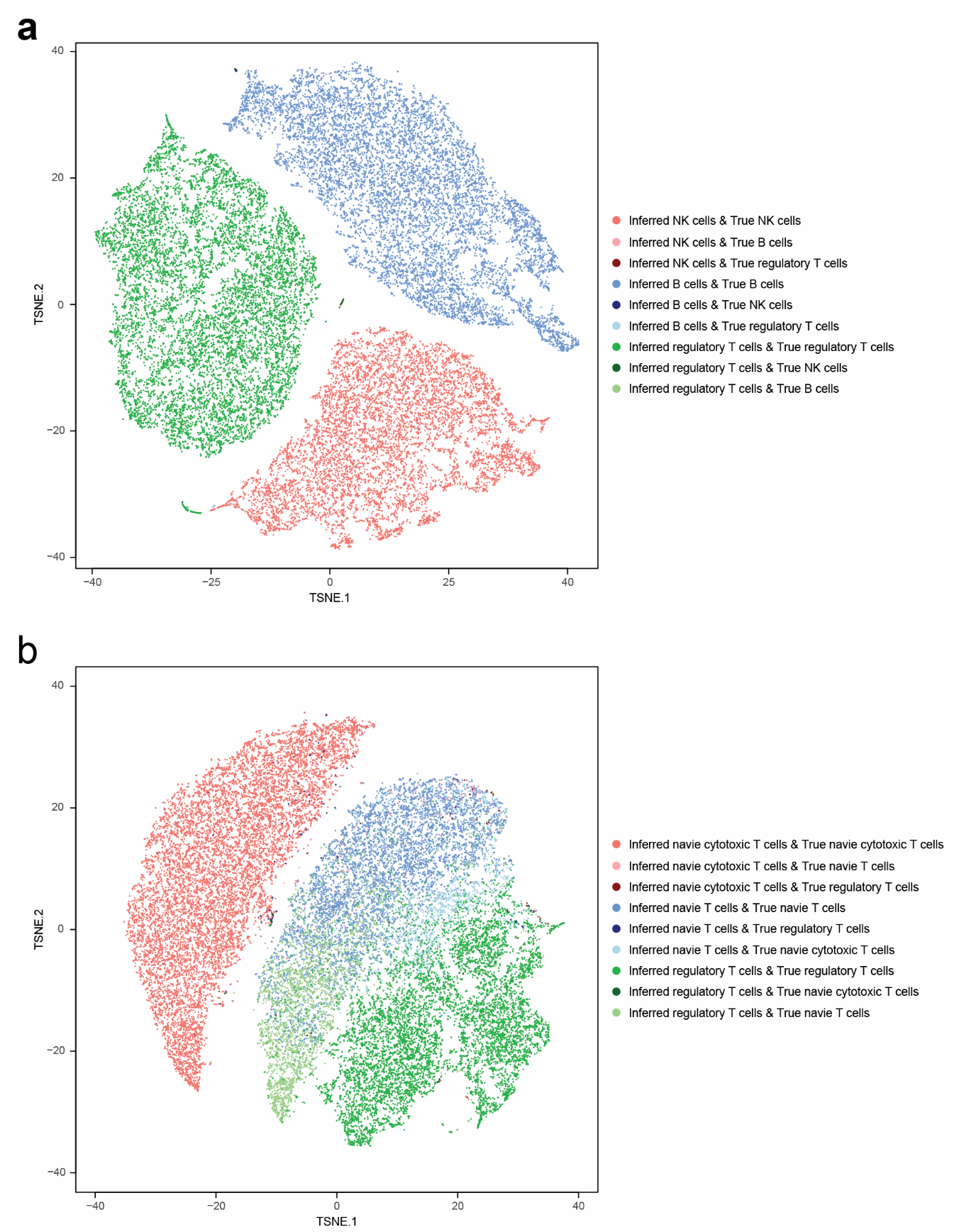

**Supplementary Fig. S2.** t-SNE visualization of comparing SAFE-clustering inferred cluster results and true cluster results for the PBMC simple case (**a**) and the PBMC challenging case (**b**).

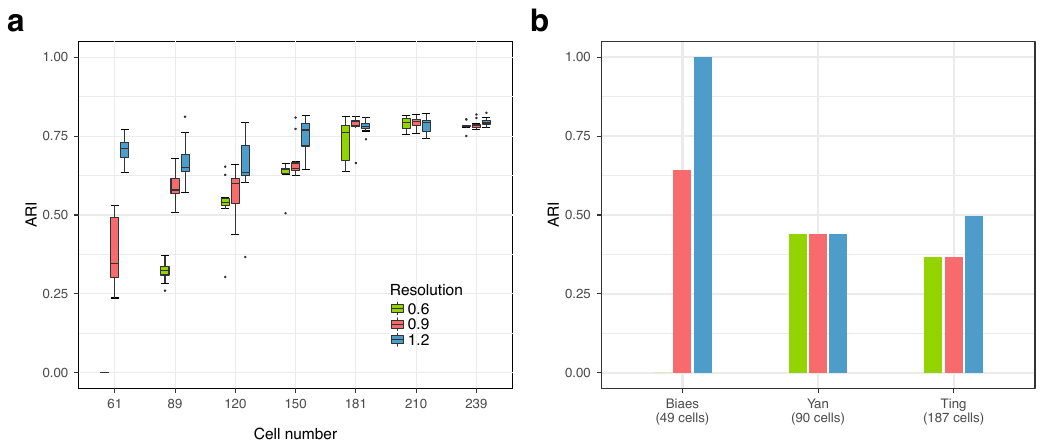

**Supplementary Fig. S3.** (**a**) Evaluation performance of Seurat with varying resolution parameters depending on sample size. (**b**) Seurat clustering performance in 3 different small datasets with varying resolution parameters.

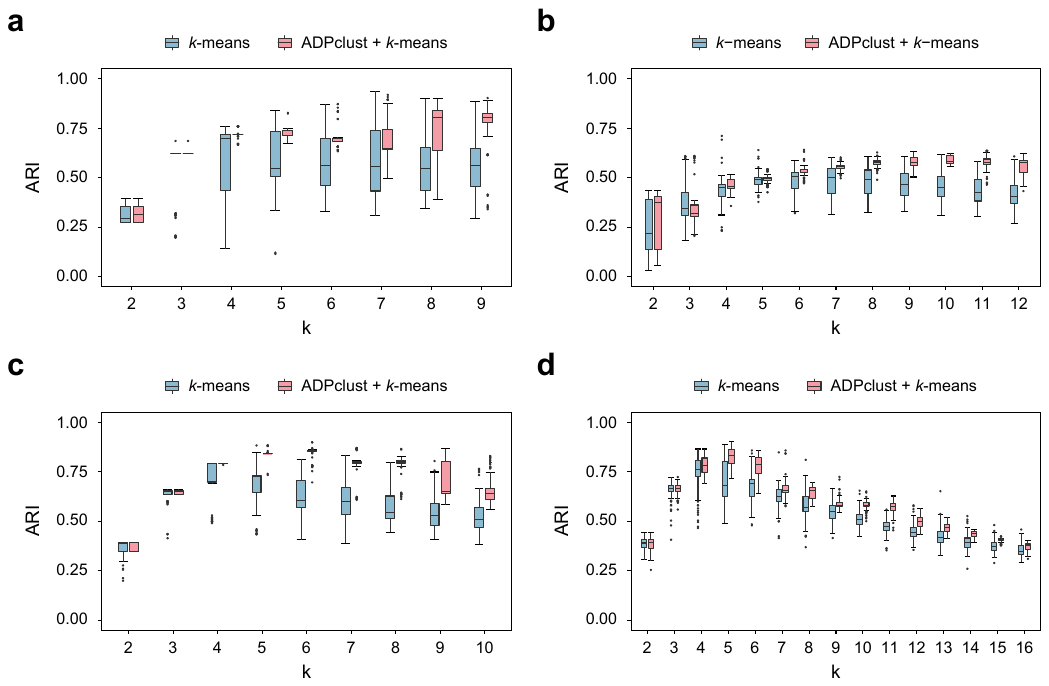

**Supplementary Fig. S4.** Comparing performance of t-SNE+kmeans with and without ADPclust centroid estimation step before kmeans clustering for four datasets (Yan, Ting, Darmanis, and Baron_human2, **a-d** respectively).

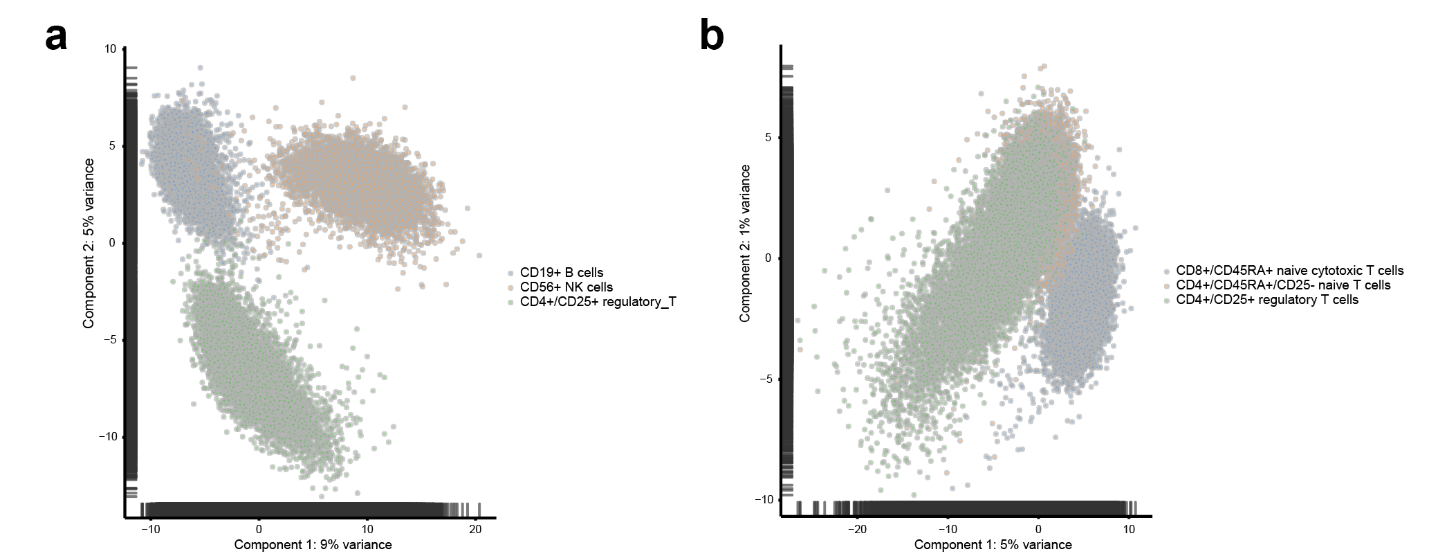

**Supplementary Fig. S5.** PCA plot for the simple case (**a**) and challenging case (**b**) to estimate the number of clusters to input for SC3.

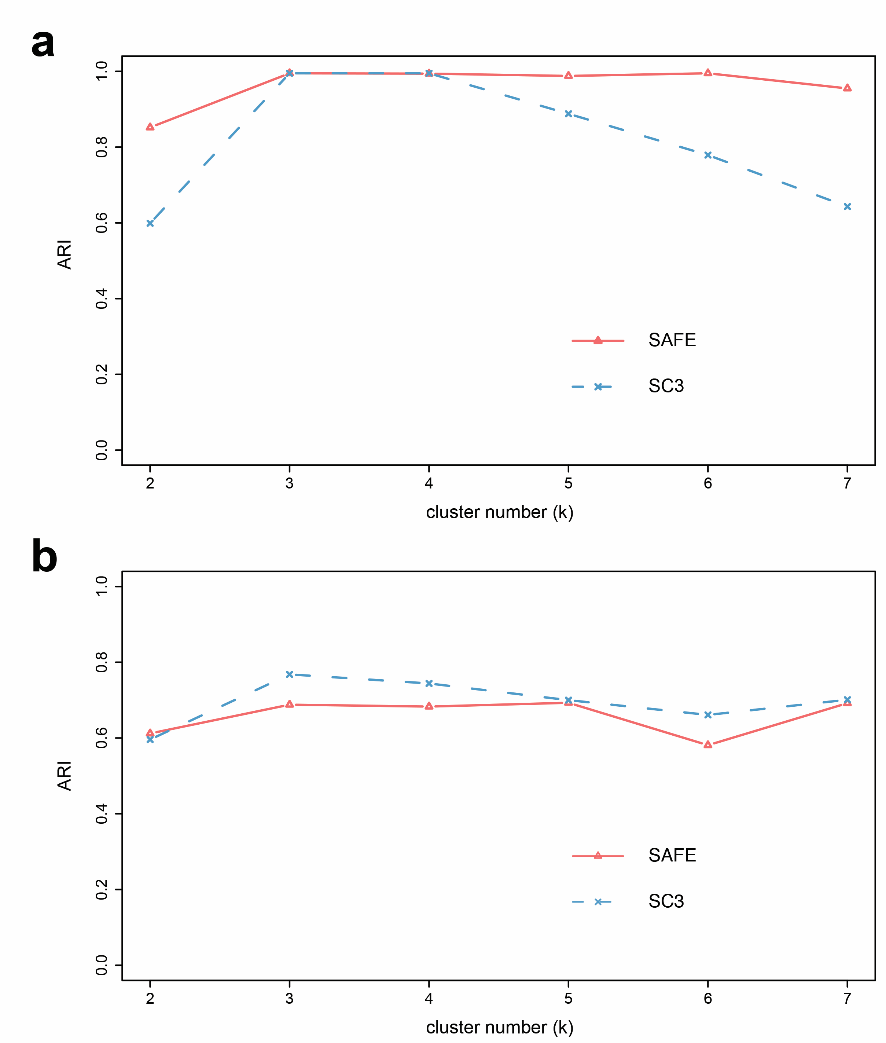

**Supplementary Fig. S6.** Assessing the robustness of SAFE-clustering results for the simple case (**a**) and challenging case (**b**) when varying the number of clusters for SC3 results and holding the other three individual methods constant.
